# Transposable element mobilization in interspecific yeast hybrids

**DOI:** 10.1101/2020.06.16.155218

**Authors:** Caiti Smukowski Heil, Kira Patterson, Angela Shang-Mei Hickey, Erica Alcantara, Maitreya J. Dunham

**Affiliations:** Department of Genome Sciences, University of Washington, Seattle, WA; Department of Biological Sciences, North Carolina State University, Raleigh, NC; National Heart, Lung, and Blood Institute, Bethesda, MD; Center for Genomics and Systems Biology, New York University, New York, NY; The New York Institute of Technology College of Osteopathic Medicine, Glen Head, NY

**Keywords:** hybridization, transposable elements, transposition rate, Ty element, Saccharomyces

## Abstract

Barbara McClintock first hypothesized that interspecific hybridization could provide a “genomic shock” that leads to the mobilization of transposable elements. This hypothesis is based on the idea that regulation of transposable element movement is potentially disrupted in hybrids. However, the handful of studies testing this hypothesis have yielded mixed results. Here, we set out to identify if hybridization can increase transposition rate and facilitate colonization of transposable elements in *Saccharomyces cerevisiae x Saccharomyces uvarum* interspecific yeast hybrids. *S. cerevisiae* have a small number of active long terminal repeat (LTR) retrotransposons (Ty elements), while their distant relative *S. uvarum* have lost the Ty elements active in *S. cerevisiae*. While the regulation system of Ty elements is known in *S. cerevisiae*, it is unclear how Ty elements are regulated in other *Saccharomyces* species, and what mechanisms contributed to the loss of most classes of Ty elements in *S. uvarum*. Therefore, we first assessed whether transposable elements could insert in the *S. uvarum* sub-genome of a *S. cerevisiae* x *S. uvarum* hybrid. We induced transposition to occur in these hybrids and developed a sequencing technique to show that Ty elements insert readily and non-randomly in the *S. uvarum* genome. We then used an *in vivo* reporter construct to directly measure transposition rate in hybrids, demonstrating that hybridization itself does not alter rate of mobilization. However, we surprisingly show that species-specific mitochondrial inheritance can change transposition rate by an order of magnitude. Overall, our results provide evidence that hybridization can facilitate the introduction of transposable elements across species boundaries and alter transposition via mitochondrial transmission, but that this does not lead to unrestrained proliferation of transposable elements suggested by the genomic shock theory.

## Introduction

Transposable elements (TEs) are mobile, repetitive genetic elements that have colonized nearly every organism across the tree of life. TEs self-encode machinery to either replicate or excise themselves from one genomic location and re-insert at another genomic location, which can disrupt genes or gene expression and promote chromosomal rearrangements through ectopic recombination. Due to the high potential of fitness costs of these mutations, most organisms have evolved host defense systems to regulate TEs (Rebollo *et al.* 2012). However, while experiments and population genetics show that the average effect of TE insertions is deleterious, individual transposition events may be neutral or even advantageous (Wilke *et al.* 1992; González and Petrov 2009; Stoebel and Dorman 2010; Van’t Hof *et al.* 2016; Hope *et al.* 2017; Li *et al.* 2018; Esnault *et al.* 2019; Niu *et al.* 2019). Far from their historical status of “junk DNA,” TEs are now known to contribute to a variety of processes including telomere maintenance (Pardue and DeBaryshe 2011), centromere structure (Casola *et al.* 2008; Carbone *et al.* 2012; Gao *et al.* 2015; Kursel and Malik 2016; Jangam *et al.* 2017), sex chromosome evolution (Bachtrog 2003) (Ellison and Bachtrog 2013; Dechaud *et al.* 2019), regulation of gene expression, evolution of genome size, karyotype, and genomic organization across the tree of life (Petrov 2002; Jiang *et al.* 2004; Gregory and Johnston 2008; Pellicer *et al.* 2014; Schubert and Vu 2016; Kapusta *et al.* 2017; Thybert *et al.* 2018; Bourque *et al.* 2018).

The type and number of TEs in a genome vary between populations and species, as do the regulatory systems organisms use to suppress TEs (Bourque *et al.* 2018). In her Nobel prize lecture in 1983, Barbara McClintock hypothesized that hybridization between different populations or species could act as a “genomic shock” that initiates TE mobilization that could lead to the formation of new species.

> “Undoubtedly, new species can arise quite suddenly as the aftermath of accidental hybridizations between two species belonging to different genera. All evidence suggests that genomic modifications of some type would accompany formation of such new species. Some modifications may be slight and involve little more than reassortments of repetitious DNAs, about which we know so little… Major genome restructuring most certainly accompanied formation of some species. Studies of genomes of many different species and genera indicate this. Appreciation of the various degrees of reassortment of components of a genome, that appear during and following various types of genome shock, allows degrees of freedom in considering such origins. It is difficult to resist concluding that some specific “shock” was responsible for the origins of new species in the two instances to be described below. The organization of chromosomes in many closely related species may resemble one another at the light microscope level. Only genetic and molecular analyses would detect those differences in their genomes that could distinguish them as species. In some instances of this type, distinctions relate to the assortment of repetitious DNAs over the genome, as if a response to shock had initiated mobilities of these elements(McClintock 1984).”

This idea revolves in part around the idea that hybridization could cause a de-repression of TE regulation, perhaps by mismatch of the repression system in the hybrid genome. Evidence supporting this hypothesis is mixed. Initial excitement centered on the hybrid dysgenesis system in *Drosophila melanogaster*, where an intraspecific cross between a strain carrying the P-element transposon to a strain without P-elements produced sterile offspring (Kidwell *et al.* 1977; Bingham *et al.* 1982; Kidwell 1983; Rose and Doolittle 1983; Bucheton *et al.* 1984). However, attempts to test this model of transposon induced speciation across other species of *Drosophila* demonstrated this applied in certain crosses but not others (Coyne 1985, 1986, 1989; Hey 1988; Lozovskaya *et al.* 1990; Labrador *et al.* 1999; Kelleher *et al.* 2012). Studies in the *Arabidopsis* species complex are similarly mixed, with evidence that crosses between *Arabidopsis thaliana* and *Arabidopsis arenosa* lead to an upregulation of the retrotransposon ATHILA, the level of which is linked to hybrid inviability (Josefsson *et al.* 2006); but crosses between *A. thaliana* and *A. lyrata* show no change in expression of TEs in interspecific hybrids (Göbel *et al.* 2018). Iconic studies in desert sunflowers revealed that three independent hybrid species formed by crosses of *Helianthus annuus* and *Helianthus petiolaris* had elevated copy number of LTR retrotransposons compared to their parent species (Ungerer *et al.* 2006, 2009; Staton *et al.* 2009). However, contemporary crosses of the same *Helianthus* parental species did not lead to large scale proliferation of TEs, although the TEs remain transcriptionally active (Kawakami *et al.* 2011; Ungerer and Kawakami 2013; Renaut *et al.* 2014). From all of these studies, there is evidence that hybridization in some cases can lead to a misregulation of the TE repression system and potential proliferation of TEs, but it remains unclear how widespread this phenomenon is and what factors contribute to this process.

In this study, we use *Saccharomyces cerevisiae x Saccharomyces uvarum* interspecific hybrids as a system to explore the hypotheses that hybridization can lead to an increase in transposition of TEs, and that hybridization could provide an avenue for colonization of a naïve genome by TEs. *S. cerevisiae* has been used as a model to understand retrotransposition for decades. *Saccharomyces* TEs are made up of Long Terminal Repeat (LTR) retrotransposons which fall into six families, Ty1, Ty2, Ty3, Ty3_1p, Ty4, and Ty5 (Kim *et al.* 1998; Carr *et al.* 2012). Ty elements make up a small fraction of the genome (<5%), with a total of approximately 50 full length Ty elements and over 400 solo LTRs in the *S. cerevisiae* reference genome (Kim *et al.* 1998; Carr *et al.* 2012). Ty1 is the most abundant and well-studied Ty element, representing almost 70% of the full length TEs in the reference genome, with its closely related family Ty2 making up a further 25%. Ty1 preferentially integrates near genes transcribed by RNA Polymerase III through an association between integrase and Pol III-complexes (Mularoni *et al.* 2012). The other families are rare; Ty3 is thought to be an active family (Hansen and Sandmeyer 1990), Ty4 has full length elements but has not been observed to transpose (Hug and Feldmann 1996), and no intact copies of Ty5 are known (Voytas and Boeke 1992).

Ty content and copy number vary across strains and species (Liti *et al.* 2005; Bleykasten-Grosshans *et al.* 2013), with Ty elements inherited vertically and horizontally (Liti *et al.* 2005; Carr *et al.* 2012; Bergman 2018; Czaja *et al.* 2020), and certain Ty families lost. For example*, S. uvarum*, a cold-tolerant species 20 million years divergent from *S. cerevisiae*, has no full length Ty elements with the exception of the Ty4-like Tsu4 (which likely evolved from the Ty4/Tsu4 superfamily which gave rise to the Ty4 element in the *S. cerevisiae*/*S. paradoxus* lineage) (Neuvéglise *et al.* 2002; Liti *et al.* 2005; Bergman 2018). While there are no intact copies of Ty1 elements in *S. uvarum*, there are a number of Ty1 and Ty2 solo LTRs, indicative of past retrotransposition events (Scannell *et al.* 2011).

*Saccharomyces* are particularly interesting because the clade has recently lost RNAi regulation of transposable elements (Drinnenberg *et al.* 2009). Instead, *S. cerevisiae* Ty1 is regulated through a novel mechanism, copy number control (CNC) (Garfinkel *et al.* 2003, 2016; Saha *et al.* 2015; Ahn *et al.* 2017). A truncated form of the Ty-encoded Gag capsid protein (p22) disrupts virus-like particle assembly in a dose-dependent manner, allowing high levels of retrotransposition when few Ty1 elements are present and inhibiting transposition as copy number increases (Garfinkel *et al.* 2005; Saha *et al.* 2015). However, re-introducing the proteins Dicer and Argonaute of *Naumovozyma castellii* to *S. cerevisiae* can restore RNAi, and are sufficient to silence endogenous Ty retrotransposition (Drinnenberg *et al.* 2009). *S. uvarum* and some strains of its close relative *S. eubayanus* are the only *Saccharomyces* species to still retain Dicer (Wolfe *et al.* 2015), but how this may contribute to Ty regulation is unclear. CNC is not well understood for Ty elements besides Ty1, nor is it known how CNC functions in other species of *Saccharomyces* outside of *S. cerevisiae* and S*. paradoxus* (Moore *et al.* 2004; Czaja *et al.* 2020).

Here, we use Ty-specific sequencing and transposition assays in lab-created interspecific hybrids to understand how hybridization impacts Ty mobilization. We show that hybridization does not lead to an increase in transposition rate or proliferation of Ty1 elements in hybrids. However, we do document variation in transposition rate in hybrids that is mediated through a curious phenomenon of mitochondrial inheritance, such that hybrids with *S. uvarum* mitochondria have a lower rate of transposition than hybrids with *S. cerevisiae* mitochondria.

## Materials & Methods

### Strains and plasmids used

Strains YMD119 and YMD120 are haploid *S. cerevisiae* strains of GRF167 background (YMD119, L35 - 102 C1, *ura3-167 MATα*, YMD120, L47-102 C1, *ura3-167 MATα*). YMD119 is a high-Ty strain created by repeated induced transposition of Ty1, while YMD120 has a Ty1 profile similar to S288C (Scheifele *et al.* 2009). These strains were crossed to YMD366, a *S. uvarum* lab strain of background CBS7001, to create hybrids YMD130, and YMD129, respectively. Strains YMD3375 (*his3d200 ura3-167*, Ty1his3AI-242 (chrXII)) and YMD3376 (*his3d200 ura3-167*, Ty1his3AI-273 (chrII)) carry an integrated, marked Ty1 element for use in transposition assays (gifts from Mary Bryk, see (Bryk *et al.* 1997)). YMD3375 and YMD3376 were crossed to CSH143 to create *S. cerevisiae* diploids CSH144 and CSH145, and to CSH189 to create *S. cerevisiae x S. uvarum* hybrids (CSH192, 193, 195-198) for transposition assays. Strains YMD3375 and YMD3376 were provided by Chris Hittinger, and were also crossed to CSH187 to create hybrids with a *S. uvarum dcr1* knockout for transposition assays. Strains yCSH215 and yCSH216 are ρ^0^ versions of YMD3375 and YMD3376, respectively, which were created via passage on ethidium bromide. The *Ty1his3AI* plasmid was a gift from David Garfinkel, as used in (Curcio and Garfinkel 1991). See **Table S1** for a list of all strains used.

### Survey of *S. uvarum* Ty elements

We downloaded sequencing reads for 54 *S. uvarum* isolates (Almeida *et al.* 2014) and aligned each sample with bwa aln (Li and Durbin 2009) to a reference genome made of Ty1, Ty2, Ty3, Ty4, and Ty5 full length elements. We then employed RetroSeq version 1.41 (Keane *et al.* 2013) on a subset of these samples to call novel insertions in the *S. uvarum* genome. Each call was manually inspected using Integrative Genomics Viewer (Robinson *et al.* 2011).

### TySeq library creation and sequencing

DNA was extracted using the Hoffman-Winston protocol (Hoffman and Winston 1987), cleaned using the Zymo Clean and Concentrate kit (Zymo Research, Irving, CA), and quantified on the Qubit fluorometer. To identify Ty elements, we took a sequencing based approach modified from previous methods (van Opijnen *et al.* 2009; Mularoni *et al.* 2012), which we call TySeq. The library preparation was based off of previously described methods (Wetmore *et al.* 2015; Sanchez *et al.* 2019), modified as described here (**Figure S1**, see **Supplemental Text** for detailed protocol, **Table S2** for primers). 1 μg of genomic DNA was sheared to an average size of 800 bp using a Covaris machine with default settings. The sheared DNA fragments were blunt ended, and A-tails were added to the fragments to ligate the Illumina adapter sequences. We used a nested PCR approach, in which we first attempted to amplify full-length Ty1 and Ty2 elements using custom primers designed to target sequences interior to Ty1 and Ty2 elements, avoiding the LTR sequences (See **Table S2** for primers used), and custom indexed primers that target the Illumina adapter sequence were used to enrich for genomic DNA with Ty1 and Ty2 insertion sites. The second PCR used the product from PCR#1 with the same indexed primer that binds the Illumina adapter, and a second primer that binds the Ty1 and Ty2 LTR and adds the second Illumina adapter (**Figure S1**). The resulting libraries were quantified on a Qubit and run on a 6%TBE gel to assess library size. Libraries were sequenced on an Illumina NextSeq 500 using a custom R1 sequencing primer that binds the Ty1 and Ty2 LTR. Due to the low complexity of the libraries, libraries were never allowed to exceed 10-15% of a sequencing run.

TySeq of induced transposition with the marked Ty1 was produced as above, except using a primer that binds to *HIS3* instead of Ty1 (See **Table S2** for primers used). Strain CSH153 was transformed with the *Ty1his3AI* plasmid and crossed to *S. uvarum* strain CSH6 to create strain CSH177. Biological replicates of CSH177 were grown overnight in C-URA media to maintain the plasmid, then a small number of cells were used to inoculate 48 replicates of 1 mL C-URA + 2% galactose, which was grown for 2 days at 20°C. Replicates were then pooled together and plated on C-HIS plates. Plates were scraped and pooled together to be used for DNA library preparation.

### TySeq sequencing analysis

Due to the sequencing primer design, Read 1 sequencing reads should start with 27bp of the LTR. We took a stringent approach to filtering TySeq reads for alignment. First, R1 reads were cropped to 27bp in length using trimmomatic v0.32 (Bolger *et al.* 2014) and aligned to a Ty element reference genome, which contained all annotated LTR and Ty elements in the *S. cerevisiae* S288C reference genome (obtained from SGD, last updated 2015-01-13), using bwa aln (Li and Durbin 2009). Only reads mapping to this Ty reference genome were used in later steps. We subset all 150 bp reads to only reads that mapped to the Ty reference genome using seqtk subseq (https://github.com/lh3/seqtk). These full length R1 reads then had the first 27 bp cropped using trimmomatic to remove the LTR specific sequence from the read. A second filtering step was taken to remove all reads mapping to Ty elements using the same approach as above. Finally, reads not mapping to Ty elements were aligned to the reference genome, sacCer3 or Sbay.ultrascaf (Scannell *et al.* 2011). Only positions with more than 50 reads were considered likely insertions. All potential inserts were visually inspected using Integrative Genomics Viewer (Robinson *et al.* 2011) and we confirmed a subset of the insertions using PCR. Genome coverage in 25 bp intervals was assessed using igvtools count (Robinson *et al.* 2011). Overlap of Ty elements between different samples was assessed using bedtools “window,” and proximity to sequence features was assessed using bedtools “closest” (Quinlan and Hall 2010).

### Transposition rate assays

Transposition rate was measured in strains with an integrated Ty1 tester *Ty1his3AI* as has been previously described (Curcio and Garfinkel 1991; Bryk *et al.* 1997; Dunham *et al.* 2015). A strain was grown overnight, then cell count was assessed by hemacytometer. Approximately 2500 cells were diluted in 10 mL of YPD then inoculated in 100 μL volume in a 96 well plate, such that there were less than 500 cells per well. The plate was sealed with a breathable membrane and incubated without shaking at 20°C for 4 days. All exterior wells were discarded. C-HIS plates were prepared for the assay by drying via blotting with sterile Watson filter paper or incubation in a 30 incubator for 2 days. Three wells were titered on YPD plates to assess population size and the remaining wells entire contents were individually, independently spotted onto very dry C-HIS plates and left to incubate at 30°C for 3 days. Patches were scored as zero or non-zero. Each assay examined on average 57 patches, with at least two biological replicates. Transposition rate was scored via a maximum likelihood method (Lea and Coulson 1949).

### Whole genome sequencing of selected hybrids

Based on results from transposition assays, four strains were selected for whole genome sequencing (yCSH195, yCSH198, yCSH193, yCSH196). Strains were grown up overnight, and a portion of each was used to start new transposition assays. The remaining cells had DNA extracted using the Hoffman Winston protocol followed by library preparation using the Illumina Nextera library kit. The samples were sequenced on an Illumina NextSeq 500 and reads were aligned to a concatenated reference genome of *S. cerevisiae* and *S. uvarum* (Scannell *et al.* 2011) using bwa mem and default parameters (Li and Durbin 2009). Read depth was assessed using igvtools (Robinson *et al.* 2011) and normalized to account for average genome wide coverage. Read depth per homolog was used as a proxy of copy number change in the hybrid.

### Plate reader assay

We used a BioTek Synergy H1 plate reader to assay growth rate by measuring OD600 every 15 minutes at 25°C with agitation over the course of 60 hours. 3 replicates of each strain (CSH218, 219, 221, 222, 224, 225, 227, 228) were grown in rich media (YPD), and 3 replicates of each strain were grown in media with glycerol as the sole carbon source (YPG).

### Data availability

Sequencing data are deposited under BioProject ID PRJNA639117.

## Results

### Nearly all isolates of *S. uvarum* are free of Ty elements

Characterization of the CBS7001 lab strain of *S. uvarum* determined that *S. uvarum* was devoid of full length Ty elements with the exception of Tsu4 (Bon *et al.* 2000; Neuvéglise *et al.* 2002; Liti *et al.* 2005; Scannell *et al.* 2011). We conducted a bioinformatics based survey of 54 worldwide isolates from natural and fermentation conditions (Almeida *et al.* 2014) to identify if the characterization of CBS7001 was representative of the species as a whole. We largely confirm *S. uvarum* to be missing full length Ty elements, but find a single strain (GM14) with a potential full length Ty1 element. This strain was isolated from grape must in France and has introgression derived from *S. eubayanus*, *S. kudriavzevii*, and *S. cerevisiae*, although the potential insertion is not in one of these regions. Given the strain’s history of hybridization, we sought to identify if hybridization could provide a possible mechanism for Ty elements to insert in naïve species’ genomes.

### TySeq, a sequencing method for detecting *de novo* transposable element insertions

Detecting TEs in sequencing data is notoriously difficult. Their repetitive nature and large size (for example, the Ty1 is approximately 6kb) present major challenges to genome assembly, and traditional alignment pipelines will miss new insertions due to their absence in the reference genome. There have been many advances in the computational detection of TEs using short read sequencing data (Ewing 2015; Rishishwar *et al.* 2017), and long-read sequencing will likely represent the new gold standard for TE annotation (Disdero and Filée 2017; Bergman 2018; Kutter *et al.* 2018; Shahid and Slotkin 2020). However, there is still a wide range of false positives and false negatives associated with computational methods, and long-read sequencing is currently more expensive and less high-throughput than short read methods. We therefore present a method, TySeq, adapted from previous methods (van Opijnen *et al.* 2009; Mularoni *et al.* 2012), which can identify novel or non-reference Ty1 element insertions. While we apply this to Ty1 and Ty2 elements in *Saccharomyces* specifically, it is easily adapted to support the detection of other TEs in other organisms.

Briefly, we created a sequencing library quite similar to traditional whole genome sequencing library methods with small modifications (**Figure S1**). We started with a sheared genomic library of 800bp, large enough to span the LTR region of Ty elements and capture flanking genomic sequence. We created a biased library by using primers that amplify DNA fragments which contain a full length Ty1 or Ty2 element. We then used a custom sequencing primer that sequences off the LTR, capturing the flanking genomic region. These reads can be mapped back to a reference genome, thus identifying locations of new, non-reference, and reference TE insertions.

We applied TySeq to *S. cerevisiae* x *S. uvarum* hybrid strains to demonstrate proof of principle (**Figure 1, Figures S2-S4**). We identified 52 putative Ty1 and Ty2 elements (read depth of 50+ reads supporting, **Table S3**) in the *S. cerevisiae* sub-genome of a wild-type hybrid strain. While the strain background differs from the *S. cerevisiae* reference genome, we find a similar number of Ty1 and Ty2 elements present. We additionally utilized a “high-Ty” hybrid, in which the *S. cerevisiae* portion of the genome carries a higher load of Ty1 elements derived from repeated induction of transposition using a synthetic construct (Scheifele *et al.* 2009). We identified 71 putative Ty1 and Ty2 elements (read depth of 50+ reads supporting, **Table S3**) in the *S. cerevisiae* sub-genome of this high-Ty hybrid. We then created a synthetic mixed population (90% wild-type hybrid, 10% high-Ty hybrid) to test the sensitivity of our TySeq protocol in detecting low frequency Ty insertions. We detected 87 Ty1 and Ty2 elements in the synthetic mixed sample, largely recapitulating Ty elements derived from both the wild-type hybrid (49/52 elements detected at a read depth of 50+ reads) and high-Ty hybrid (69/72 elements detected at read depth of 50+ reads), indicating we can detect most Ty elements which are only present in 10% of a population. The overall false positive rate (detected in the mixed sample but not in either wild-type nor high-Ty strain) is 8/87 (or 9.20%) and the false negative rate is 5/87 (5.75%, present in wild-type and/or high-Ty strain but not in mixed sample). The majority of both false positive and false negative detected insertions are the result of presence of an element with 50 or more reads in one sample, with reads between 1-49 read depth in the other sample(s) (**Table S3**). We did not identify Ty1 or Ty2 elements in the *S. uvarum* sub-genome of these hybrid strains, consistent with the previously identified absence of full length Ty1 or Ty2 elements in *S. uvarum* (**Figure 1, Figures S2-S4**). This furthermore suggests that new insertions do not occur in the outgrowth of the colony from a single hybrid zygote.

**Figure 1:**
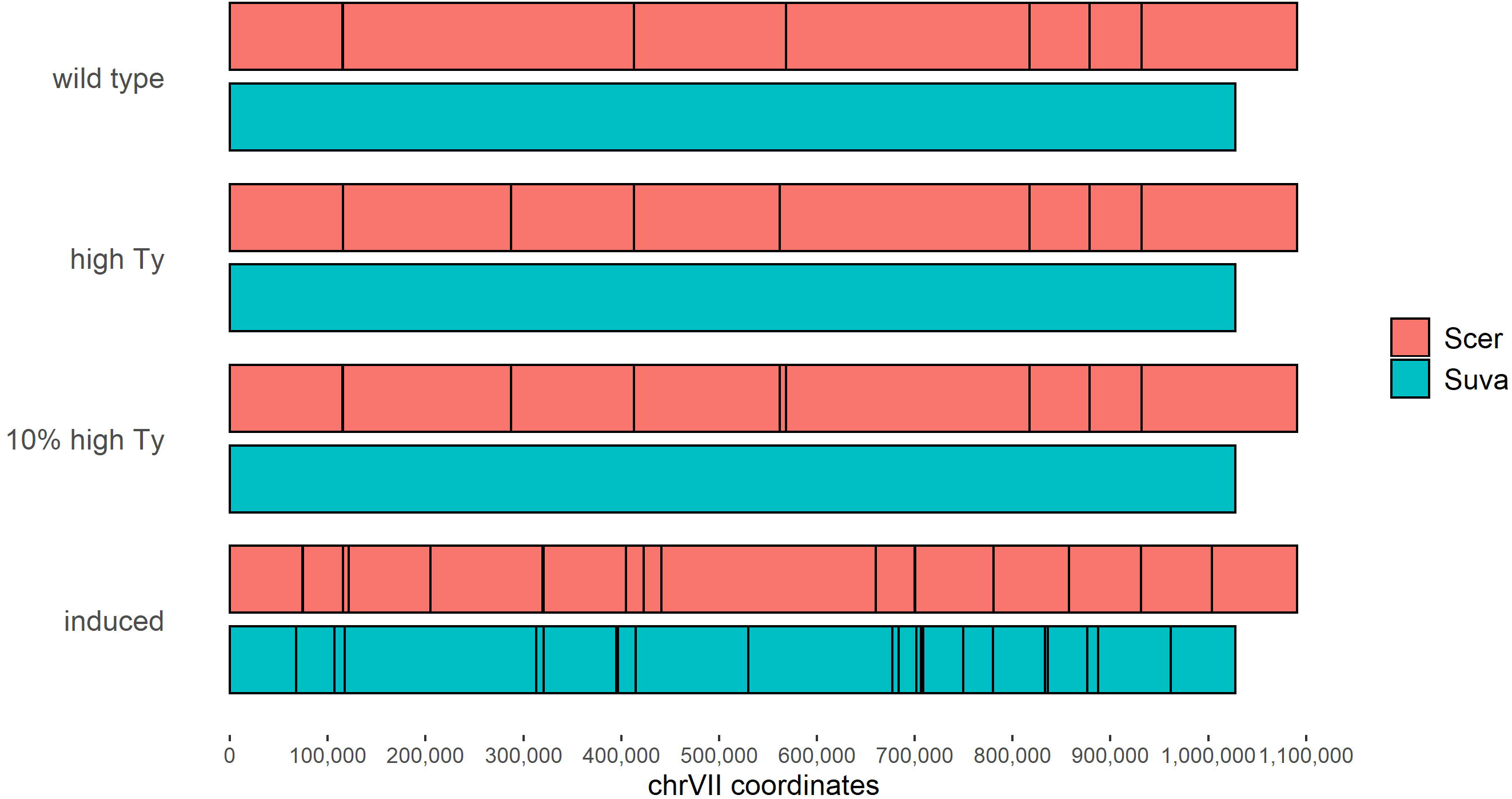
Using TySeq to identify Ty elements in *S. cerevisiae* x *S. uvarum* hybrids. Ty elements detected with TySeq are shown as black lines across chrVII for the *S. cerevisiae* (pink) and *S. uvarum* (blue) portions of a hybrid genome. Ty elements are shown for wild type (YMD129), high-Ty (YMD130), a mixed sample of 90% YMD129 and 10% YMD130, and a pool of His+ colonies obtained from induced transposition. No Ty elements were detected in the *S. uvarum* portion of the hybrid genome except when transposition was artificially induced (these insertions are plotted using *S. uvarum* genome coordinates). For whole genome figures, see **Figures S2-S4,S6-S7.** For coordinates of insertions, see **Tables S3-S5**.

We next sought to identify if we could induce transposition and detect novel insertions in a hybrid genome, and in particular, if insertions would occur in the *S. uvarum* sub-genome. We used a marked Ty1 element, *Ty1his3AI* on a plasmid under galactose induced expression(Curcio and Garfinkel 1991). This construct has a full length Ty1 element with a *HIS3* reporter gene interrupted with an artificial intron. Upon transposition, the intron is spliced out, restoring functionality to *HIS3* and allowing detection of transposition events by growth on media lacking histidine (**Figure S5**). We sequenced two replicates of a pool of His+ colonies and detected 23,693 and 31,083 reads mapping to the *S. cerevisiae* sub-genome, and 33,427 and 45,272 reads mapping to the *S. uvarum* sub-genome. We identified 93 and 122 insertions in the *S. cerevisiae* sub-genome respectively (with 50+ reads, **Table S4, Figure 1, Figures S6, S7**), with many of these sites differing from those identified in the wild-type and high-Ty hybrid. A similar number of insertions were identified in the *S. uvarum* sub-genome, with 121 and 109 insertions detected respectively (**Figure 1, Figures S6, S7, Table S5**). These results suggest that Ty1 is equally likely to insert into either *S. cerevisiae* or *S. uvarum* genomes.

In *S. cerevisiae,* Ty1 elements preferentially insert near PolIII transcribed genes, like tRNAs (Mularoni *et al.* 2012). Here, we show that in the two replicates, 83.68% and 88.55% of reads that map to the *S. uvarum* genome are within 2kb of an annotated tRNA gene. This is similar to the 93.6% reported for *S. cerevisiae* (Mularoni *et al.* 2012), suggesting the insertion preference for Ty1 is conserved despite 20 million years divergence between the two species. The discrepancy between *S. cerevisiae* and *S. uvarum* might be due in part to differences in annotation between the two species reference genomes (there are fewer tRNA genes annotated in the *S. uvarum* reference). Our results thus show that Ty1 elements can insert in the *S. uvarum* genome, and suggest that hybridization may be a mechanism through which transposable elements could hop from one species to another.

### Variable transposition rate in hybrids

We then directly measured transposition rate in *S. cerevisiae x S. uvarum* hybrids to test the hypothesis that transposition is increased in interspecific hybrids. We used *S. cerevisiae* strains which have a marked Ty1 element, *Ty1his3AI*, integrated on chrII and chrXII, respectively (**Table S1, Figure S5**). These marked *S. cerevisiae* strains were crossed to an unmarked *S. cerevisiae* strain to create diploids, and to an unmarked *S. uvarum* strain to make hybrids. Transposition rate was scored via the fluctuation method (Lea and Coulson 1949).

Transposition rate is dependent on the location of the marked Ty1 element, and can depend upon ploidy, where diploids may have a lower rate of transposition compared to haploids due to MATa/α repression (Elder *et al.* 1981; Herskowitz 1988; Garfinkel *et al.* 2005). We first repeated transposition assays in marked *S. cerevisiae* haploids and recapitulate previously published results, that *S. cerevisiae* haploid *Ty1his3AI* strains have transposition rates of 10^−6^ to 10^−7^ per generation (Curcio and Garfinkel 1991, 1992; Bryk *et al.* 1997). We furthermore recapitulate results of similar haploid and diploid rates (**Table 1**) (Garfinkel *et al.* 2005 p. 1).

**Table 1:**
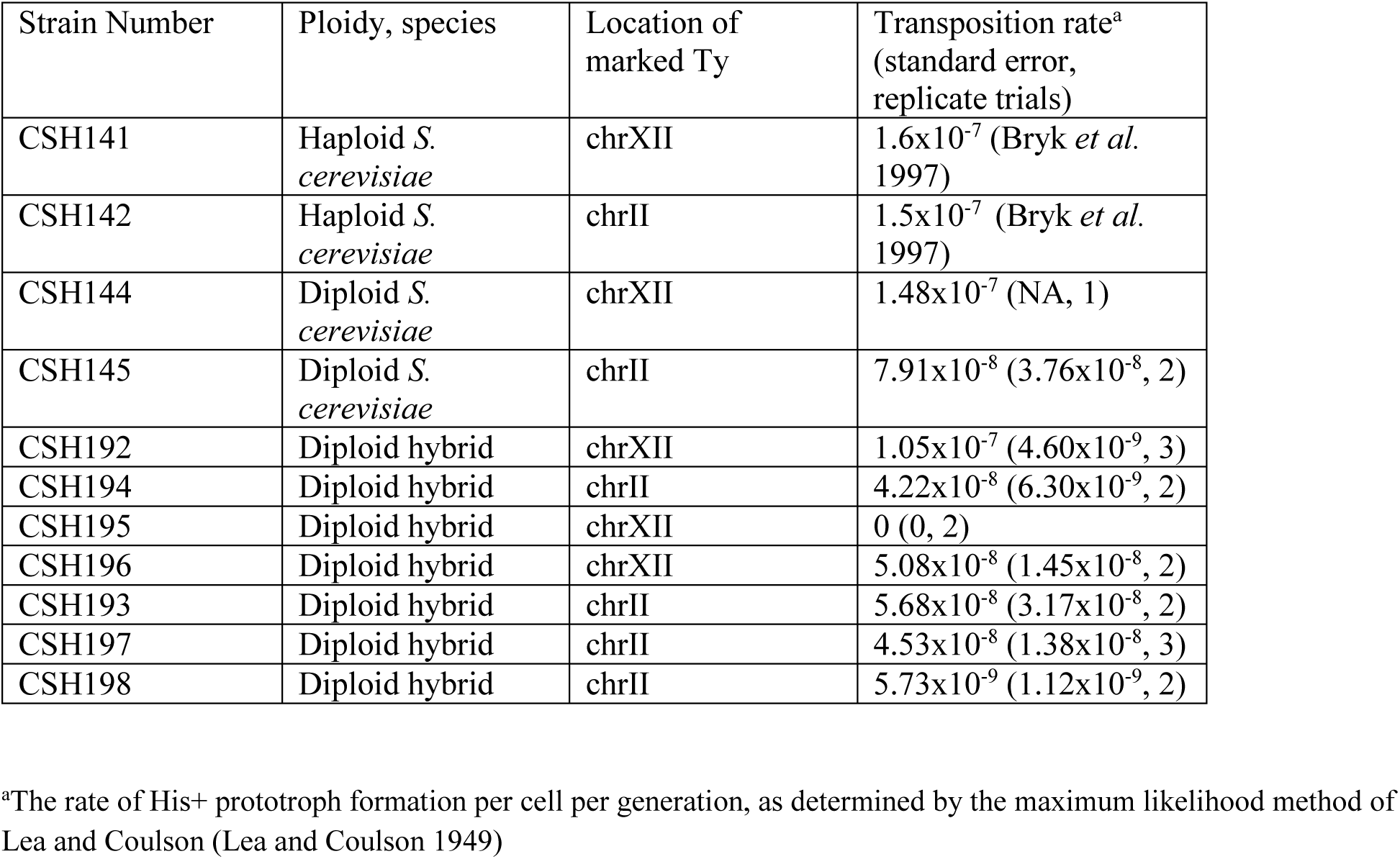
Variable transposition rate across hybrids.

We tested the hypothesis that the maintenance of one of the RNAi genes, Dicer (*DCR1*), in *S. uvarum* may be responsible for the absence of most Ty elements in that species. *DCR1* is absent in *S. cerevisiae*, so hybrids would normally have only the single *S. uvarum* copy of *DCR1*. We created a hybrid with a *S. uvarum dcr1* knockout. If *DCR1* mediates transposition rate, we would expect that *dcr1* hybrids would have an increased transposition rate. Instead, we found the rate in these hybrids to be 5.44 × 10^−8^ (+/− 5.26 × 10^−9^), similar to the rate observed in hybrids with an intact copy of *S. uvarum DCR1* (**Table 1**).

We tested transposition rate in 7 independent hybrid crosses (**Table 1**). We clearly show that hybridization does not increase transposition rate, with the highest rate of transposition observed in hybrids at approximately 1.05 × 10^−7^ (+/− 4.60 × 10^−9^), similar to rates in haploid *S. cerevisiae*, ranging to undetectably low levels of transposition (scored as a rate of 0). This variation in transposition rate between hybrids is significant (p=0.0049, ANOVA). Hybrids should be isogenic within a cross, and between crosses should only be differentiated by the marked Ty1 element residing on chrII or chrXII. Differences in transposition rate between independent hybrid matings could result from copy number variation resulting from genomic instability following hybridization, a point mutation or insertion/deletion that occurred during the growup of the culture for the transposition assay, or differential mitochondrial inheritance.

To identify the causal variants contributing to transposition rate variation in these hybrids, we selected strains that exhibited a low transposition rate (yCSH195, yCSH198), and strains with a diploid-like transposition rate (yCSH193, yCSH196) for whole genome sequencing. We identified a loss of the *S. cerevisiae* copy of chrXII in yCSH195, which resulted in the loss of the marked Ty1, hence the observed rate of 0 (**Figure S8**). We did not identify any other copy number variants, point mutations, or insertion/deletions in the remaining strains; however, we observed that the other hybrid with low transposition rate (yCSH198) inherited the *S. uvarum* mitochondrial genome (mtDNA), while the other strains (yCSH193, yCSH196) inherited the *S. cerevisiae* mtDNA. mtDNA is inherited from one parent (uniparental inheritance) in almost all sexual eukaryotes (Birky 1995, 2001), including the *Saccharomyces* yeasts. Previous work has observed a transmission bias in *S. cerevisiae x S. uvarum* hybrids, which typically inherit the *S. cerevisiae* mtDNA, although there are a variety of genetic and environmental factors that contribute to mtDNA inheritance such as temperature and carbon source (Marinoni *et al.* 1999; Lee *et al.* 2008; Hsu and Chou 2017; Hewitt *et al.* 2020). Mitotype can affect a number of phenotypes, such as temperature tolerance in yeast hybrids (Baker *et al.* 2019; Li *et al.* 2019; Hewitt *et al.* 2020), but to our knowledge has not been previously implicated in transposition.

### *S. uvarum* mtDNA decreases transposition rate in *S. cerevisiae x S. uvarum* hybrids

We set out to test the hypothesis that mitotype can influence transposition rate in hybrids by creating a set of crosses with controlled mtDNA inheritance. We induced strains of *S. cerevisiae* and *S. uvarum* to lose their mtDNA (denoted as ρ^0^) through passage on ethidium bromide, then crossed these ρ^0^ strains to the corresponding species with mtDNA intact. We conducted transposition assays in these newly created hybrids and demonstrate that the inheritance of *S. uvarum* mtDNA results in a significantly lower transposition rate (p=0.0039, Welch’s t-test; **Table 2**). A series of growth curves on fermentable and non-fermentable carbon sources illustrates that *S. uvarum* mtDNA is still functioning in respiration, although results in a slightly slower growth rate than the identical strain with *S. cerevisiae* mtDNA (**Figure S9**).

**Table 2:** *S.uvarum* mtDNA decreases hybrid transposition rate by an order of magnitude.

| Strain Number | Ploidy, species | mtDNA | Transposition rate<sup>a</sup> (standard error, replicate trials) |
| --- | --- | --- | --- |
| CSH218 | Diploid hybrid | *S. uvarum* | 0 (0,2) |
| CSH221 | Diploid hybrid | *S. uvarum* | 5.28x10<sup>-9</sup> (1.09x10<sup>-9</sup>, 3) |
| CSH224 | Diploid hybrid | *S. cerevisiae* | 6.51x10<sup>-8</sup> (1.44x10<sup>-8</sup>, 3) |
| CSH225 | Diploid hybrid | *S. cerevisiae* | 3.97x10<sup>-8</sup> (1.09x10<sup>-8</sup>, 3) |

<sup>a</sup>The rate of His<sup>+</sup> prototroph formation per cell per generation, as determined by the maximum likelihood method of Lea and Coulson (Lea and Coulson 1949)

## Discussion

In summary, we combined a modified sequencing strategy, TySeq, with *in vivo* transposition rate assays to test the hypothesis that TE mobilization may be increased in interspecific hybrids. Using an integrated, marked Ty element construct to quantify transposition rate, we identified significant variation in transposition rate among strains that we expected to be isogenic. We show that mitochondrial inheritance can explain this variation, with *S. uvarum* mtDNA decreasing transposition rate in hybrids by an order of magnitude. Thus, while we reject the hypothesis that hybridization increases TE mobilization, we demonstrate hybridization can impact transposition rate in novel ways.

### Intrinsic and extrinsic variables that affect transposable element movement

There is considerable variation in TE content across species and between populations, and many extrinsic and intrinsic factors that mediate transposition rate. Both the rate and distribution of TEs are governed by their overall deleterious effect (Charlesworth and Langley 1989). All organisms have evolved defenses to limit TE movement, although these systems vary across species and include zinc-finger proteins, small RNA-based silencing strategies, DNA methylation, and chromatin modifications (Rebollo *et al.* 2012). TE elements and their host defense systems continue to evolve, which in turn changes transposition rate. For example, Kofler *et al.* utilized experimental evolution to observe the evolution of a P-element invasion in populations of naïve *D. simulans*, documenting the emergence over time of P-element specific piRNAs that curbed the spread of the P-element (Kofler *et al.* 2018). In *S. cerevisiae* and *S. paradoxus*, recent work discovered two variants of the Ty1 element segregating in populations of wild and human-associated strains that determine rates of Ty mobility (Czaja *et al.* 2020). Strains with the canonical Ty1 element show reduced mobility of canonical Ty1 whereas strains with the divergent Ty1’ (and lack of genomic canonical Ty1) show increased mobility of canonical Ty1. This is a result of the TE defense system (CNC) being Ty specific, such that Ty1’ CNC cannot control the mobility of Ty1.

Here, we find that mitochondrial inheritance in hybrids significantly changes transposition rate, the first study to document this connection. A mechanism of how mtDNA is influencing transposition is unclear, although mitochondria function in a huge variety of processes beyond generating cellular energy (Malina *et al.* 2018; Dujon 2020; Hose *et al.* 2020). The unique pattern of mtDNA inheritance and large numbers of nuclear-encoded mitochondrial genes contribute to mito-nuclear incompatibilities that underlie some speciation events (Lee *et al.* 2008; Gershoni *et al.* 2009; Chou and Leu 2010; Burton and Barreto 2012; Crespi and Nosil 2013) and human diseases (Duchen and Szabadkai 2010; Vafai and Mootha 2012). Moreover, species specific inheritance of mtDNA in hybrids results in a strong environmentally dependent allele preference for one species’ alleles or the other (Hewitt *et al.* 2020). Perhaps this species specific allele expression results in the suppression of *S. cerevisiae* encoded Ty elements in a hybrid with *S. uvarum* mtDNA, causing the observed lower rates of transposition.

Temperature also seems to play a mediating role in mitochondrial inheritance, mitochondria function, and transposable element movement. Mitochondria have been repeatedly implicated in adaptation to different temperatures (e.g., the “mitochondrial climatic adaptation hypothesis”) (Mishmar *et al.* 2003; Ruiz-Pesini *et al.* 2004; Ballard and Whitlock 2004; Wallace 2007; Dowling 2014; Camus *et al.* 2017). For example, in hybrids between thermotolerant *S. cerevisiae* and cryotolerant *S. uvarum* or *S. eubayanus*, *S. cerevisiae* mtDNA confers growth at high temperatures, while *S. uvarum or S. eubayanus* mtDNA confers growth at low temperatures (Baker *et al.* 2019; Li *et al.* 2019; Hewitt *et al.* 2020). An Australian cline of *D. melanogaster* showed thermal performance associated with each mitotype corresponds with its latitudinal prevalence (Camus *et al.* 2017). Intriguingly, TEs were shown to play a significant role in adaptation to the climatic variables in this same *D. melanogaster* cline (González *et al.* 2008, 2010). Recently Kofler et al. used experimental evolution of *D. simulans* at cold and warm temperatures and showed that temperature drastically impacts the rate at which a TE can spread in a population (Kofler *et al.* 2018). In *S. cerevisiae*, rates are estimated to be 100 fold higher at temperatures 15-20°C than at the normal lab conditions of 30°C (Paquin and Williamson 1984; Garfinkel *et al.* 2005). All transposition assays were conducted at the standard 20°C in this study, but future work could explore how temperature impacts transposition rate in non *S. cerevisiae* species, particularly the cold tolerant *S. uvarum* and *S. eubayanus*. If transposition rate is increased at cold temperatures, reduced transposition rate may be an evolutionary response to curb TE mobilization in cryotolerant species. This is certainly an intriguing area for further study.

### The role of transposable elements in evolution

In recent years we have witnessed a shift from viewing TEs as solely parasitic genetic elements, to appreciating the myriad ways in which TEs impact eukaryotic evolution. In our own work in laboratory evolution experiments, we have shown that Ty elements are often breakpoints for adaptive copy number variants and that insertions can cause adaptive gain and loss of function mutations. Intriguingly, we have previously observed fewer copy number variants in *S. uvarum* than *S. cerevisiae* evolved populations, perhaps related to their paucity of repetitive elements to facilitate such mutational events (Smukowski Heil *et al.* 2017, 2019). Copy number events, and in particular chromosome rearrangements can cause inviability between crosses (e.g., chromosomal speciation), which may represent more relevant paths in which TEs may impact speciation. While the evidence that TE mobilization in hybrids can facilitate speciation is limited, there remains much to be explored regarding evolution of host-TE dynamics between closely related species.

## Acknowledgements

Thanks to Marcus Annable for help in shearing TySeq libraries. Thanks to Mary Bryk and Chris Hittinger for sharing strains, and David Garfinkel for sharing the Ty1his3AI plasmid. This work was supported by the National Science Foundation (grant number 1516330 to MJD). CSH was supported in part by the NIH/NHGRI Genome Training Grant T32 HG00035. During this work, MJD was a Senior Fellow in the Genetic Networks program at the Canadian Institute for Advanced Research and a Rita Allen Foundation Scholar. MJD is supported in part by a Faculty Scholars grant from the Howard Hughes Medical Institute.

## Supplementary Figure legends

**Figure S1: TySeq, a modified sequencing method for detecting novel transposition events. A.** A Ty1 element is a long terminal repeat (LTR) retrotransposon. It is flanked by directional repeats (light blue). **B.** Genomic DNA is sheared into ~800 bp fragments. **C.** Illumina Nextera adapters are ligated on to the fragment ends (green). **D.** First round PCR is completed using a Ty1/Ty2 specific forward primer and a reverse primer that binds to the Nextera adapter and has a unique index (purple) and the flow cell (red). **E.** A second round of PCR is done on the Ty1 and Ty2 enriched library using the same reverse primer and a forward primer that binds to the LTR immediately adjacent to genomic sequence. **F.** A unique R1 sequencing primer is added to the sequencing run. **G.** Reads are mapped back to the reference genome, identifying sites of likely transposable element insertions.

**Figure S2: Ty elements detected with TySeq in wild type hybrid strain YMD129.** Chromosomes are shown in order from the top, chrI-chrXVI, with *S. cerevisiae* chromosomes in pink and *S. uvarum* chromosomes in blue. *S. cerevisiae* genomic coordinates are used for *S. cerevisiae* chromosomes and *S. uvarum* genomic coordinates are used for *S. uvarum* chromosomes. Black lines indicate a Ty element detected with a read depth of at least 50 reads. No Ty elements were detected in the *S. uvarum* portion of the hybrid genome. See **Table S3** for Ty element coordinates and maximum read depth.

**Figure S3: Ty elements detected with TySeq in the high-Ty hybrid strain YMD130.** Chromosomes are shown in order from the top, chrI-chrXVI, with *S. cerevisiae* chromosomes in pink and *S. uvarum* chromosomes in blue. *S. cerevisiae* genomic coordinates are used for *S. cerevisiae* chromosomes and *S. uvarum* genomic coordinates are used for *S. uvarum* chromosomes. Black lines indicate a Ty element detected with a read depth of at least 50 reads. No Ty elements were detected in the *S. uvarum* portion of the hybrid genome. See **Table S3** for Ty element coordinates and maximum read depth.

**Figure S4: Ty elements detected with TySeq in a mixed sample of hybrid strains YMD130 (10%) and YMD129 (90%).** Chromosomes are shown in order from the top, chrI-chrXVI, with *S. cerevisiae* chromosomes in pink and *S. uvarum* chromosomes in blue. *S. cerevisiae* genomic coordinates are used for *S. cerevisiae* chromosomes and *S. uvarum* genomic coordinates are used for *S. uvarum* chromosomes. Black lines indicate a Ty element detected with a read depth of at least 50 reads. No Ty elements were detected in the *S. uvarum* portion of the hybrid genome. See **Table S3** for Ty element coordinates and maximum read depth.

**Figure S5: Transposition assay with a marked Ty1. A.** A full length Ty1 element with a *HIS3* reporter gene. The HIS3 gene is interrupted with an artificial intron, and the strain cannot grow on media lacking histidine. **B.** When the Ty1 element is transcribed, the intron is spliced out, restoring the function of the *HIS3* gene, **C.** which can be detected by growth on media lacking histidine (**D.**). Independent cultures with a marked Ty1 element are grown independently and then plated on media lacking histidine. Any colonies indicate transposition has occurred. Figure adapted from (Curcio and Garfinkel 1991).

**Figure S6: Ty elements detected with TySeq from induced transposition in a hybrid, replicate 1.** Chromosomes are shown in order from the top, chrI-chrXVI, with *S. cerevisiae* chromosomes in pink and *S. uvarum* chromosomes in blue. *S. cerevisiae* genomic coordinates are used for *S. cerevisiae* chromosomes and *S. uvarum* genomic coordinates are used for *S. uvarum* chromosomes. Black lines indicate a Ty element detected with a read depth of at least 50 reads. See **Table S4** and **S5** for Ty element coordinates and maximum read depth.

**Figure S7: Ty elements detected with TySeq from induced transposition in a hybrid, replicate 2.** Chromosomes are shown in order from the top, chrI-chrXVI, with *S. cerevisiae* chromosomes in pink and *S. uvarum* chromosomes in blue. *S. cerevisiae* genomic coordinates are used for *S. cerevisiae* chromosomes and *S. uvarum* genomic coordinates are used for *S. uvarum* chromosomes. Black lines indicate a Ty element detected with a read depth of at least 50 reads. See **Table S4** and **S5** for Ty element coordinates and maximum read depth.

**Figure S8: A copy number plot of two hybrid genomes.** The top panel is yCSH193, which has a diploid-like transposition rate and no copy number changes. The bottom panel is yCSH195 and lost the *S. cerevisiae* portion of chrXII which contained the marked Ty1 element, resulting in an undetectably low transposition rate. Purple denotes a region where both alleles are present at a single copy, blue denotes a *S. uvarum* change in copy number, red denotes a *S. cerevisiae* change in copy number. Note, copy number was derived from sequencing read depth at homologous ORFs.

**Figure S9:** Growth curves of hybrids with *S. cerevisiae* mtDNA (strains CSH224, CSH225, CSH227, CSH228; red) or *S. uvarum* mtDNA (CSH218, CSH219, CSH221, CSH222; blue) over 24 hours. Each strain was grown in 3 replicates per condition and averaged. Straight lines are growth in rich medium (YPD), dotted lines are growth in a non-fermentable carbon source, glycerol (YPG). Error bars reflect standard error.

