## Supplementary figures and images for "Transposable element mobilization in interspecific yeast hybrids"

### Supplementary Figures

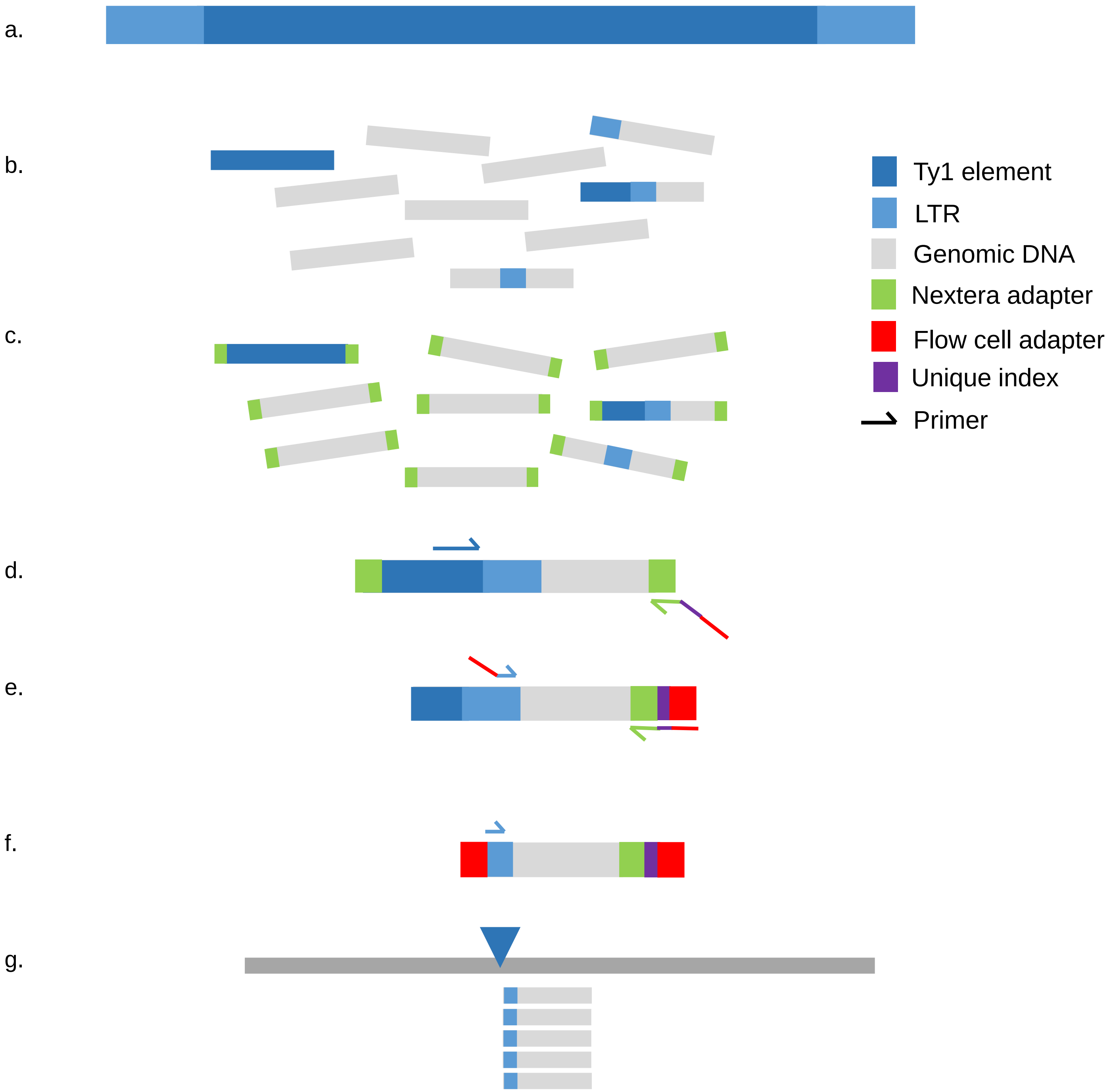

Figure S2

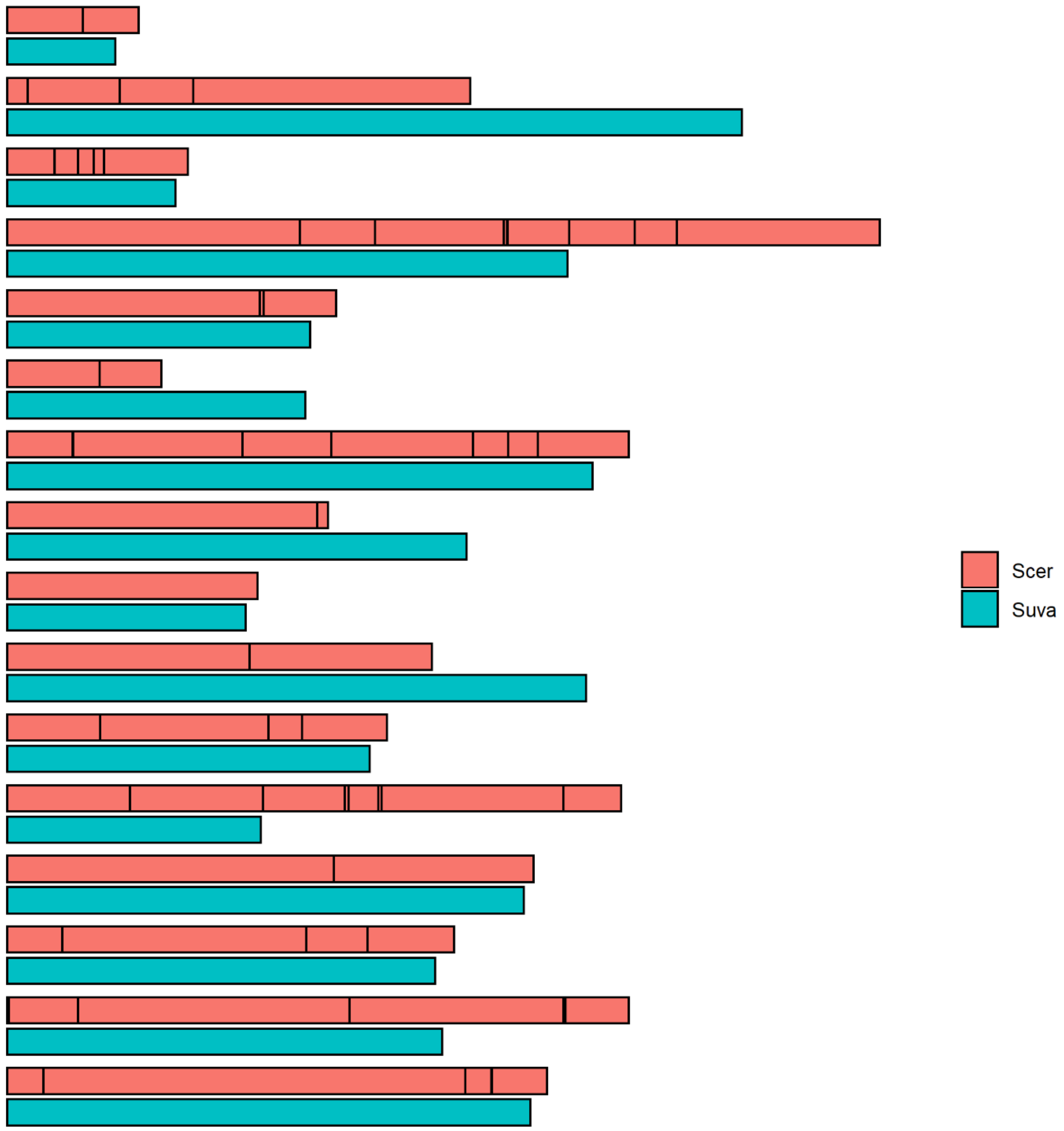

Figure S3

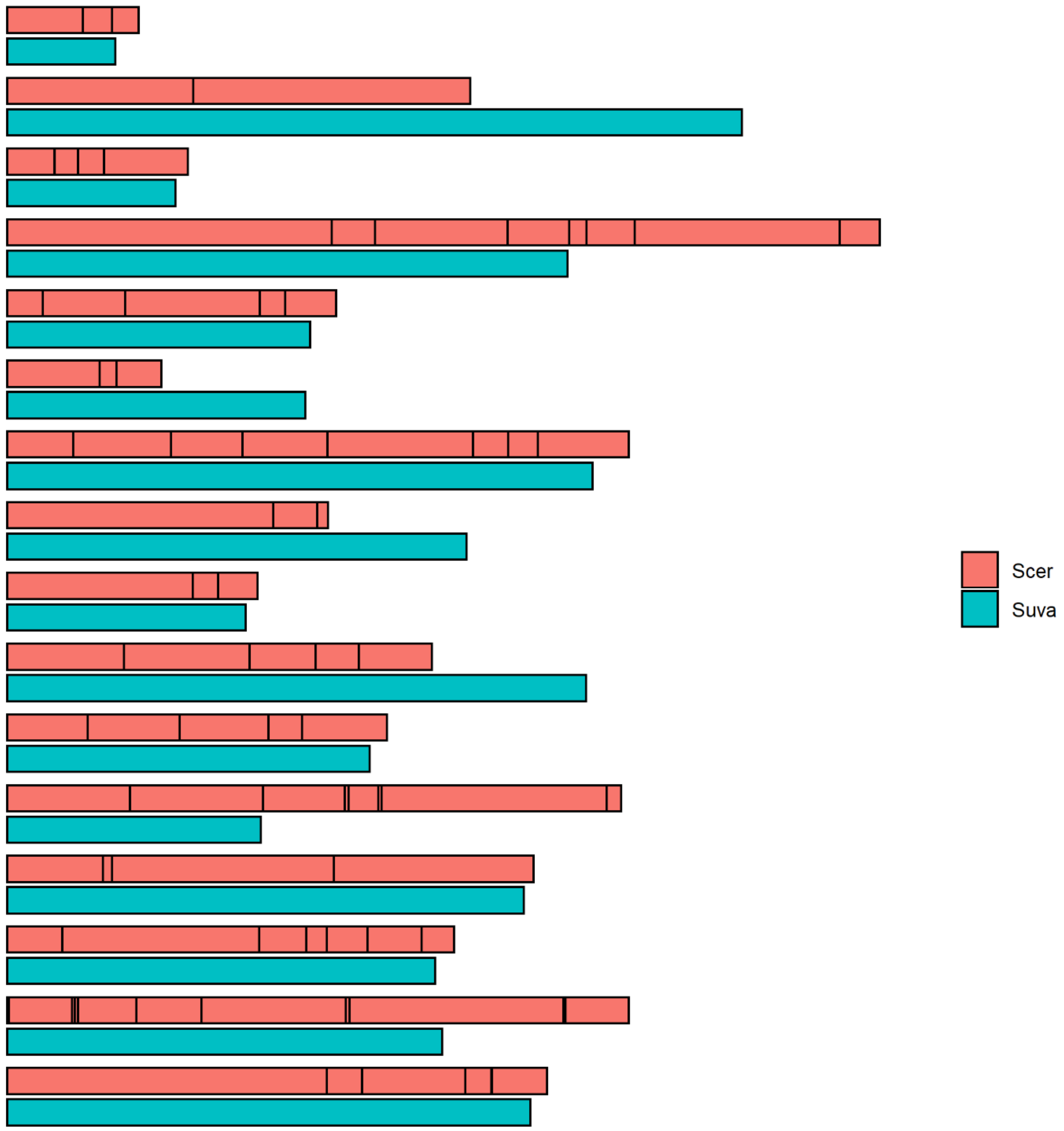

Figure S4

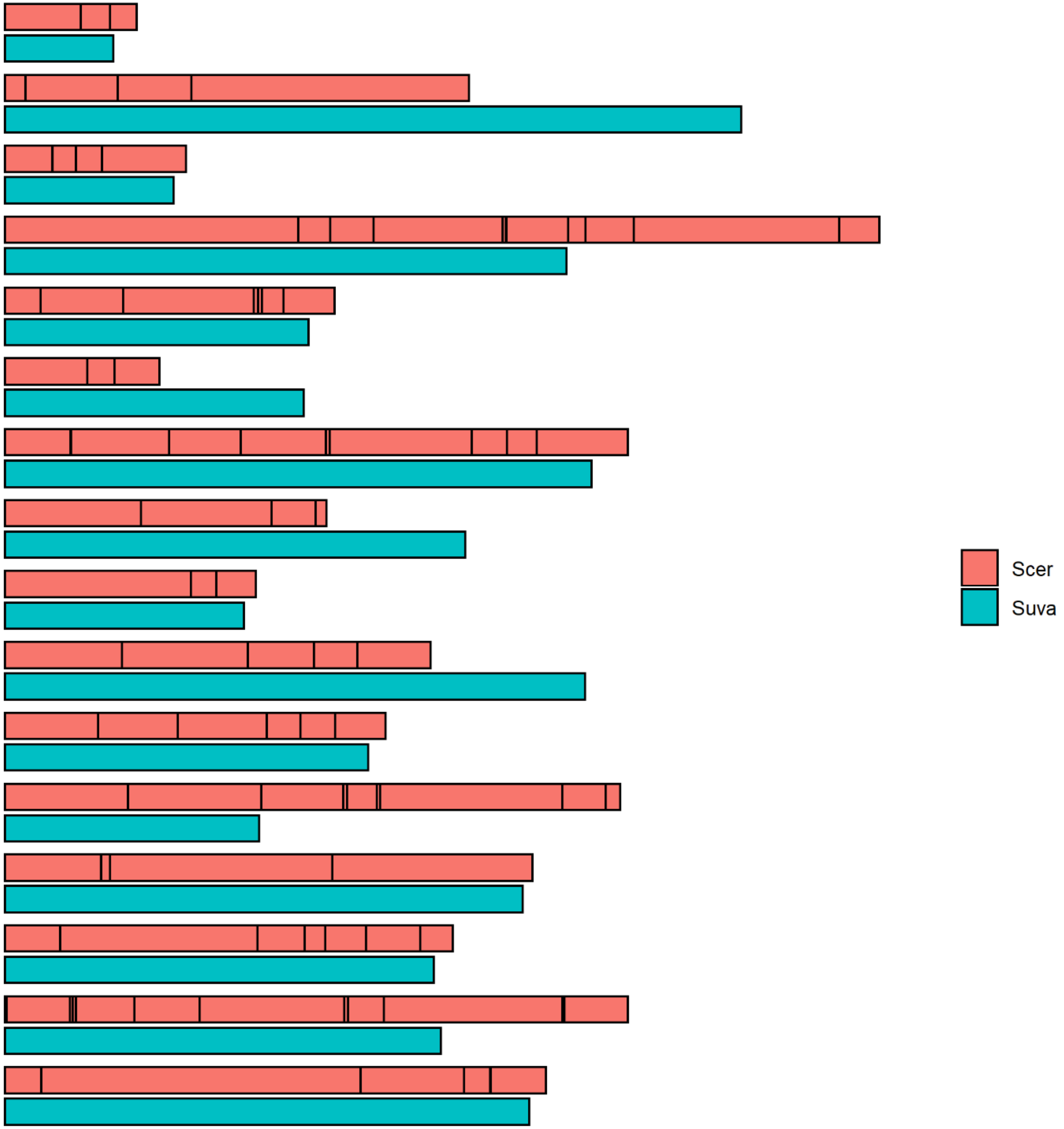

Figure S5

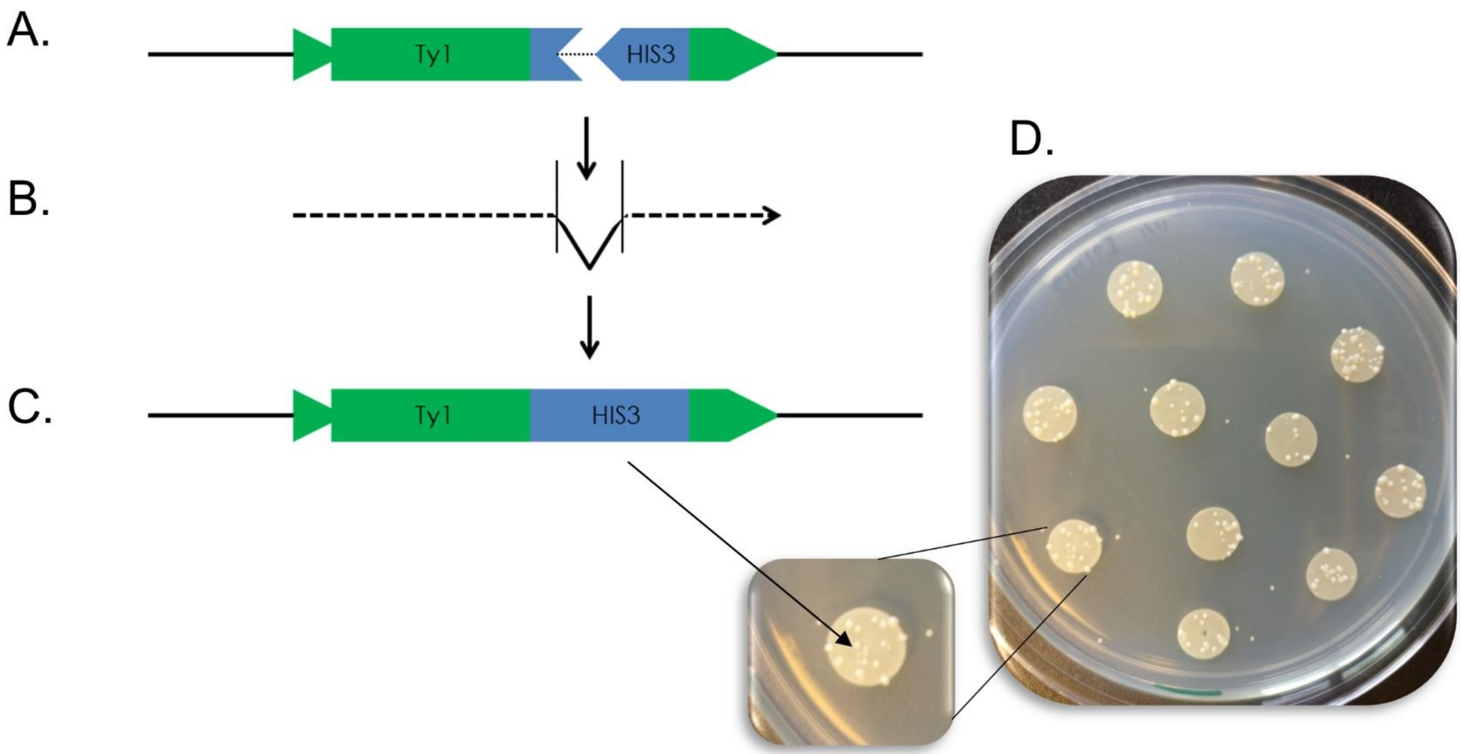

Figure S6

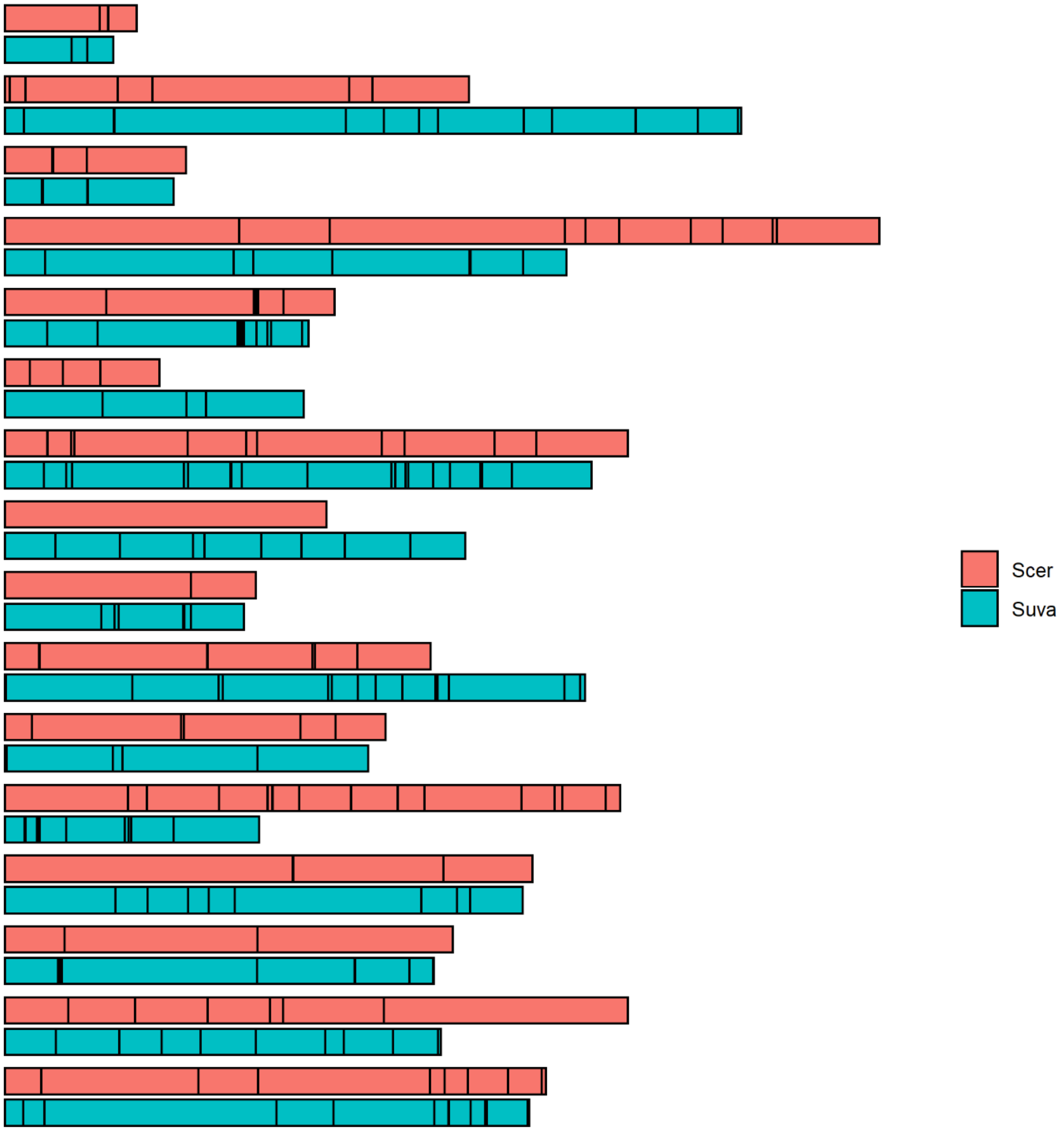

Figure S7

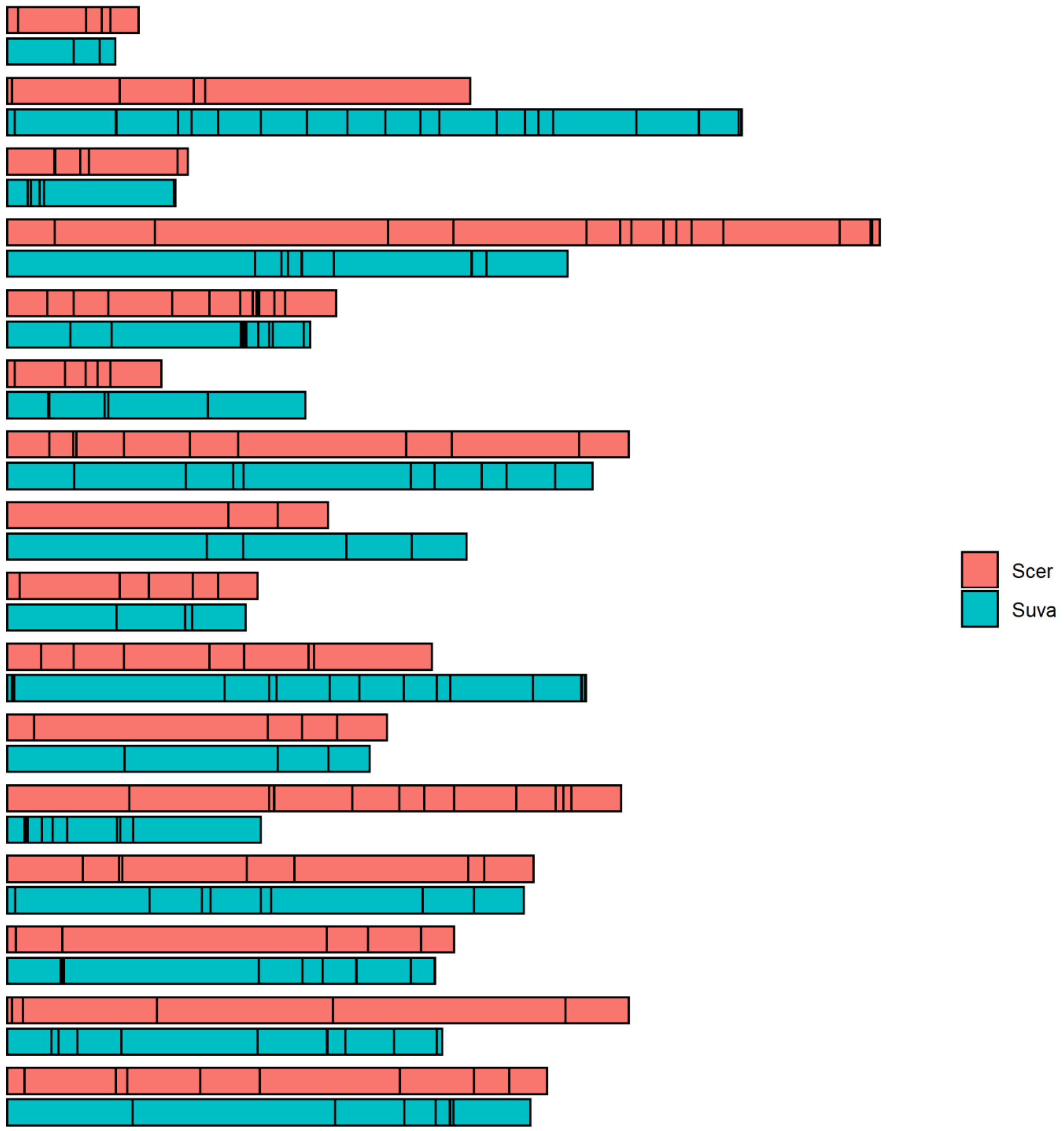

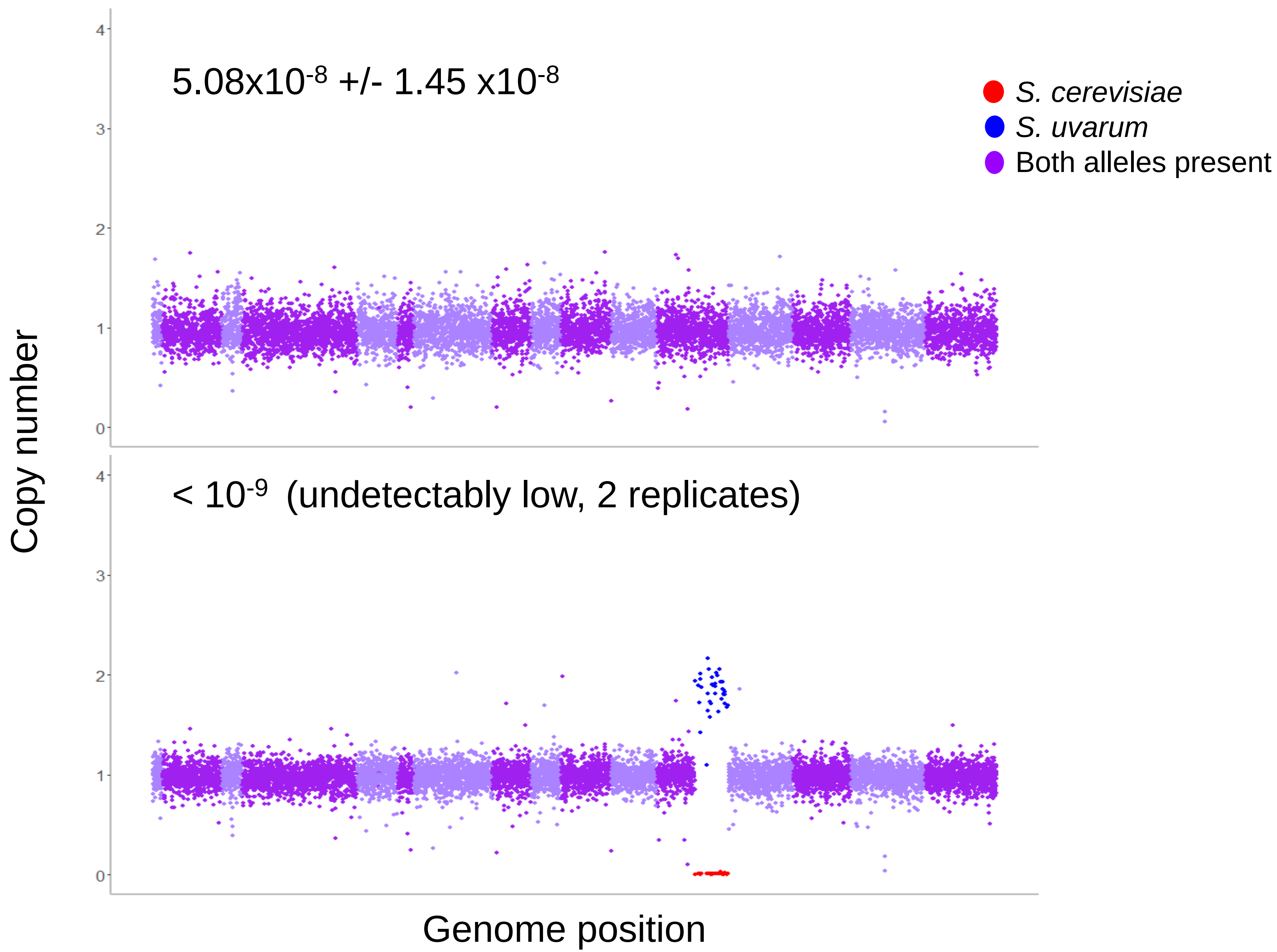

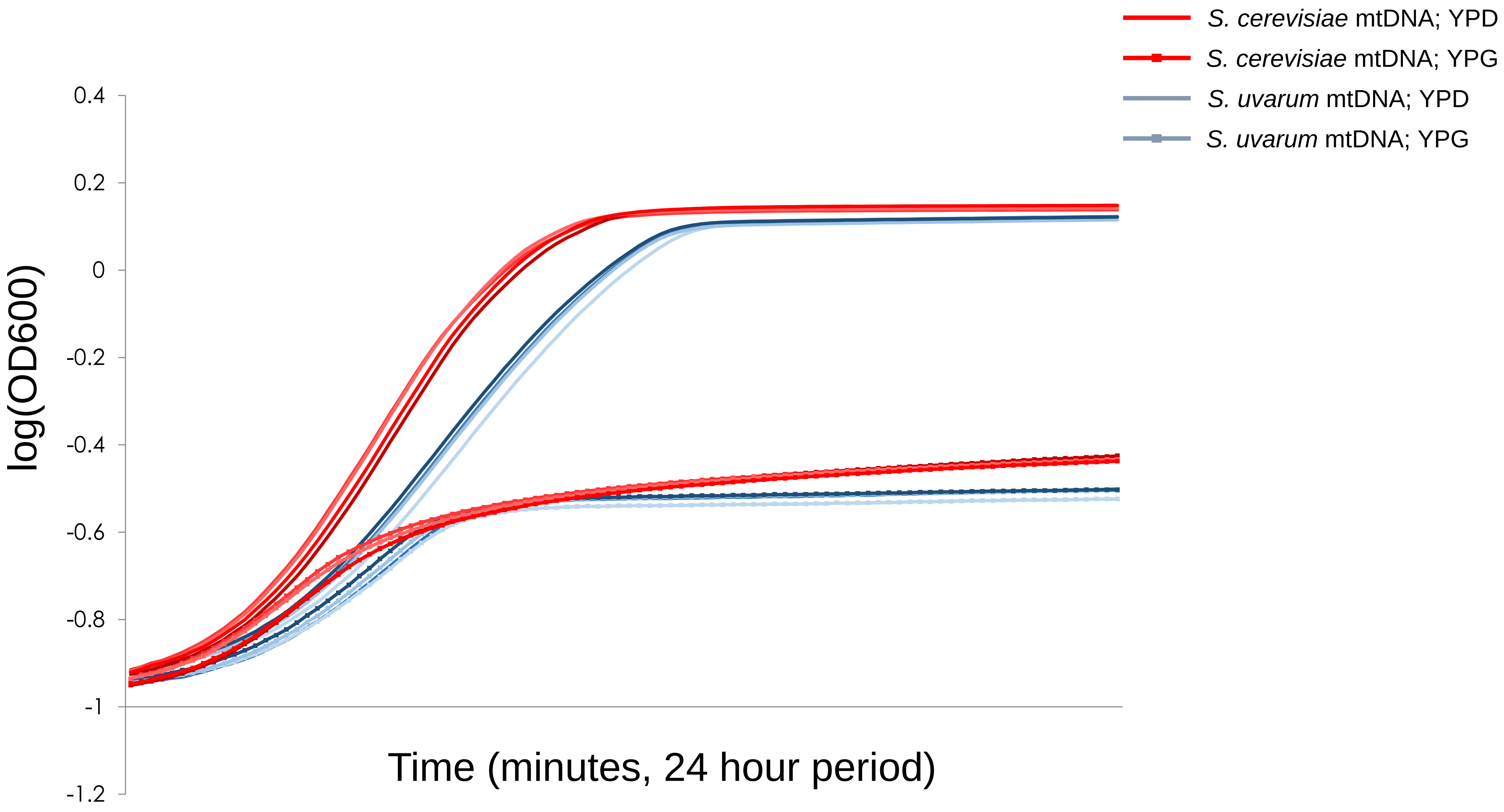
